# Analyzing Genomic Foundation Models for Viral Sequence Identification

**DOI:** 10.64898/2026.09.25.754388

**Authors:** Veronika Bůžková, Bohuslav Dvorský, Mariana Komárková, Kateřina Kollinová, Radim Krupička, Ondřej Klempíř

## Abstract

Rapid and accurate identification of viral sequences underpins clinical diagnostics, epidemiological surveillance, and the safety testing of biological products, yet the established alignment-based methods such as BLAST are inherently closed-set, with accuracy tied to how well a query is already represented in the reference database. Recent advances in genomic foundation models (GFMs) have enabled alignment-free approaches to sequence classification. However, their performance relative to traditional methods remains incompletely characterised. We evaluated GFMs, specifically the Nucleotide Transformer (NT) and DNABERT families, for viral sequence classification using a dataset derived from the Reference Viral Database (RVDB). Sequence embeddings were generated with mean, max, and CLS pooling, classified by nearest-neighbour search in an embedding index and compared with BLAST as a reference. Matching BLAST provided a particularly stringent benchmark here, because as a closed-set method it compares each query directly against the exact reference sequences it stores, whereas a foundation model must rely on representations shaped by broad, general-purpose pre-training. On sequences of standard length, the best model, NTv3-650M, reached an accuracy of 0.93, approaching the 0.97 achieved by BLAST. On long genomic sequences, evaluated on a much smaller test set, the difference narrowed further, with NTv3-650M reaching 0.95 against 0.98 for BLAST. Robustness analyses showed that all GFMs were markedly more sensitive to base masking than BLAST. Both methods indexed the same labelled reference sequences, and BLAST compared queries with them directly at the nucleotide level, so approaching it from frozen, general-purpose representations without any parameter update is a meaningful result. NTv3-650M emerged as the most promising candidate for further optimisation and deployment in bioinformatics applications.

## 1. Introduction

Viruses are among the most abundant and genetically diverse biological entities. While many are harmless, a substantial subset are major pathogens of humans, animals, and plants, and their relevance extends well beyond acute infection. Prior viral exposures have also been linked to chronic, neurodegenerative conditions in large-scale biobank analyses and in metagenomic studies of the gut virome [1-3]. Beyond neurodegeneration, several viruses are known to be oncogenic. Detecting such viruses despite their ongoing sequence variation requires classification approaches that remain robust to mutations [4]. Accurate viral detection is equally critical for the safety testing of biological products, where next-generation sequencing (NGS) is increasingly applied to screen vaccines and biotherapeutics for adventitious viral contaminants, with a corresponding need for detection pipelines that can be documented and reviewed in a regulatory setting [5]. Across clinical diagnostics, epidemiological surveillance, and product safety, the same computational problem recurs. Viral sequences have to be identified and taxonomically classified rapidly, accurately, and at scale. A range of sequence-based detection methods addresses this need, trading off speed, cost, and sensitivity [6].

NGS sequences directly and at scale, without relying on prior knowledge of what is present in a sample. This untargeted, metagenomic approach has transformed virology by enabling the discovery of both known and previously uncharacterised viruses. Realising this potential, however, depends on two things, a comprehensive, well-curated reference database, such as the Reference Viral Database (RVDB), and computational methods capable of accurately classifying reads against it [7]. Among these computational methods, alignment-based approaches, most notably BLAST, remain the established standard. They identify matches by comparing a query directly against reference sequences at the nucleotide level, not by any learned or abstracted representation. This direct comparison makes BLAST highly sensitive and well validated whenever a query has a close homolog already present in the database, which is the case for most well-characterised viruses. Its taxonomic assignment is therefore tightly coupled to the database itself [8]. In effect, BLAST behaves as a closed-set retrieval method. It excels at recognising what it has already seen, but its accuracy is a direct function of how well a query is already represented in the reference collection.

Recent advances in artificial intelligence have given rise to powerful genomic foundation models (GFMs), which offer a fundamentally different approach to sequence analysis. Built on deep learning architectures originally developed for natural language processing, these models are pre-trained on large, broadly sampled genomic corpora (spanning many species, often including viruses) and encode nucleotide sequences as dense embeddings that capture local motifs and long-range dependencies without explicit alignment [9,10]. The field has advanced rapidly from the first Nucleotide Transformer (NTv1) and the first DNABERT model to systems that now operate at the megabase scale, including Evo 2 [11], efficient generative models such as Carbon [12] that reach hundreds of kilobases at markedly lower computational cost, and the latest Nucleotide Transformer generation (NTv3).

Because these representations are learned from broad genomic corpora, they are general-purpose, and this has already been put to practical use. Tools such as ViraLM [13] and hybrid frameworks such as VIRALpre, which fuses embeddings with k-mer features [14], have shown that foundation-model representations can support viral identification.

How well such representations perform has so far been assessed mainly by general-purpose benchmarks, which evaluate models predominantly on human and functional genomics tasks and largely omit viral classification [15]. Dedicated viral benchmarks have appeared only very recently. ViroBench, for instance, evaluated dozens of nucleotide foundation models on viral taxonomy and host classification and reported that their accuracy degraded sharply under phylogenetic and temporal distribution shifts, that pre-training taxonomic diversity matters more than parameter scale, and that alignment-based BLAST still outperformed many foundation models by directly exploiting close database matches [16]. Such evaluations, however, rely on supervised classification heads, i.e., extra layers trained per task on top of the model’s embeddings. The alignment-free deployment scenario most relevant to screening, assigning taxonomy directly from frozen embeddings placed in a similarity index, remains comparatively unexplored. It is therefore unclear whether the inductive biases learned from mostly eukaryotic pre-training corpora transfer to compact, rapidly evolving viral genomes that span highly divergent families and atypical compositional biases.

An embedding is the fixed-length representation a GFM produces for a sequence, intended to capture the properties that distinguish it, or its class, from others. How faithfully it does so is highly sensitive to implementation choices that are rarely characterised together for retrieval tasks, namely the token pooling strategy (mean, max, or CLS) used to collapse per-token outputs into that single representation, dimensionality reduction for index scalability, batch effects during embedding generation, and the handling of sequences that exceed a model’s context window. The leading model families also exhibit substantial architectural heterogeneity. They differ in tokenisation strategy, positional encoding, context length, model size, and pre-training objectives. Examples include k-mer tokenisation in DNABERT, byte-pair tokenisation in DNABERT-2, single-nucleotide tokenisation in NTv3, and the species-aware contrastive objective used in DNABERT-S. It is not known which of these factors govern performance on viral sequences [10,17,18]. Robustness is a related concern. Local alignment degrades gracefully under substitutions, whereas how GFM embeddings respond to mutation, masking, and sequencing error is critical for real reads yet uncharacterised for viral identification.

We address these gaps by evaluating GFMs for viral sequence classification under a controlled, alignment-free protocol. We focus on the Nucleotide Transformer (NT) and DNABERT families because, between them, they span the tokenisation and architectural diversity outlined above, from k-mer and byte-pair to single-nucleotide tokenisation, while remaining broadly accessible and computationally tractable at the scale of nine models evaluated here, unlike substantially larger genome-scale systems such as Evo 2. Using a dataset derived from RVDB, we compared nine models from these two families, characterising the effect of pooling strategy (mean, max, and CLS), embedding dimensionality, and long-sequence handling on classification by nearest-neighbour search. Search itself is performed with FAISS, a library for efficient similarity search over large collections of embeddings [19]. We further assess robustness to base substitution and masking across a range of corruption rates on a representative subset of models, and benchmark all models against BLAST as a reference baseline.

## 2. Materials and Methods

### 2.1 Reference database

All datasets were derived from the Reference Viral DataBase (RVDB) [7], a curated collection built specifically to reduce runtime and improve the signal-to-noise ratio of virus detection in NGS data relative to large general-purpose databases. RVDB is assembled mainly from GenBank and RefSeq using a semantically refined (SEM-R) selection process that maximises viral, virus-related, and virus-like content while minimising non-viral (cellular or bacteriophage) sequences. The clustered database (C-RVDB) is derived from the unclustered one (U-RVDB) by grouping sequences that share at least 98% nucleotide identity and keeping a single representative of each group, which removes near-duplicate records. We used the latest release available at the time of the study (v31.0, 9 January 2026); the C-RVDB contained 1,291,450 records spanning 66,700 unique organisms, and was selected over the U-RVDB to avoid near-duplicate sequences. RVDB contains not only complete genomes but also complete and partial coding sequences (CDS) and defined genomic regions.

### 2.2 Dataset construction

An initial cleaning step removed sequences containing ambiguous nucleotides (the character N) or other non-standard FASTA characters, indicative of poor sequencing quality or data corruption. This excluded 409,856 sequences, roughly a third of the database. Because the benchmark subsequently samples only organisms that still retain at least 500 sequences (primary dataset) or 20 long sequences (long-sequence dataset), this step did not restrict the class coverage of the datasets used here. It also keeps ambiguous positions fully under experimental control, since N characters are introduced deliberately and at known rates in the perturbation experiments described below. Discarding a third of the records is defensible for a controlled benchmark, but it would be the wrong choice for a reference database intended for use, because a sequence with a handful of ambiguous positions is still a perfectly usable BLAST target or reference embedding. The filter applied here is therefore a property of the benchmark design and not a recommendation for how to assemble a reference database. From the cleaned database we constructed the two benchmark datasets used throughout, referred to as the primary dataset and the long-sequence dataset, using a fixed random seed for reproducibility and exporting all splits as FASTA files for downstream embedding generation and BLAST evaluation. Their composition is summarised in Table 1.

**Table 1.** Composition of datasets. The primary dataset was refined from 40 to 33 classes on the basis of the error analysis in Section 3.2, and that refinement was subsequently verified against ICTV taxonomy in Section 4. Perturbed sets are corrupted copies of the primary test set, so they share its class structure. In every dataset the training split is the indexed reference collection and the test split provides the queries.

| Property | Primary dataset | Long-sequence dataset | Perturbed sets |
| --- | --- | --- | --- |
| Source | C-RVDB v31.0 | C-RVDB v31.0 | Primary test set |
| Sequence length | up to 2,046 bp | above 50,000 bp (84,587 to 409,110 bp in the test split) | up to 2,046 bp |
| Classes | 40, refined to 33 | 16 | 33 |
| Sequences per class | 500 | at least 20 | 150 (test only) |
| Train and test split | 70/30, stratified by class | 20/80, inverted to probe low training-data availability | same split as the primary dataset |
| Training sequences | 350 per class | about 4 per class | not applicable |
| Test sequences | 150 per class (4,950 in the 33-class version) | about 16 per class (256 in total) | 4,950 per corruption setting |
| Corruption | none | none | mutation and masking, each at 0, 1, 2, 5, 10, and 20%, giving a $6 \times 6$ grid of 36 cells, of which 35 are corrupted |
| Total sequences | 20,000 (40 classes);<br>16,500 after refinement | about 320 (about 64 indexed, 256 queried) | 4,950 per corrupted cell |

#### Primary dataset

From the cleaned records, sequences longer than 2,046 bp were excluded, and only organisms retaining at least 500 sequences after this length filter were eligible. Forty classes were randomly selected, and for each, 500 sequences were sampled and split 70/30, stratified by class, into 350 training and 150 test sequences (a fixed random seed was used throughout). The sequence-length distribution was comparable between the training and test splits, ensuring that the different sequence types (complete genomes, complete and partial CDS) were equally represented in both. Following a data-quality check described in Section 3.2, which identified taxonomically ambiguous labels, a refined 33-class version of this dataset was used for all final comparisons.

#### Long-sequence dataset

Sequences exceeding 50,000 bp were retained, and classes with at least 20 such sequences were identified. Manual review identified 16 classes (including Monkeypox virus, Variola virus, African swine fever virus, White spot syndrome virus, and several herpesviruses). This dataset is drawn from the same database but covers largely different organisms from the primary dataset, because qualifying for it requires many sequences above 50,000 bp, which excludes the compact genomes that dominate the primary dataset. To additionally probe behaviour under low training-data availability, the split was inverted to about 20% training (about 4 sequences per class) and 80% test (about 16 sequences per class). Test sequences ranged from 84,587 to 409,110 bp.

#### Perturbed datasets

To simulate sequencing errors and low-quality base calls, the clean primary test set was perturbed with two independent corruption types applied at randomly selected positions, i.e. random single-nucleotide substitutions (mutations) and replacement of bases with the ambiguous character N (masking). Each was applied at six rates (0, 1, 2, 5, 10, and 20% of sequence length), and all combinations were evaluated, giving 6 × 6 = 36 test files. The training database was kept clean (0% mutation, 0% masking) throughout.

### 2.3 Models

We evaluated nine GFMs from the NT and DNABERT families: NTv1 (Multispecies 2.5B), NTv2-50M, NTv2-250M, NTv2-500M, NTv3-8M, NTv3-100M, NTv3-650M, DNABERT-2, and DNABERT-S. DNABERT and NTv1/NTv2 are BERT-style transformer encoders [20], whereas NTv3 uses a U-Net-like architecture that enables single-nucleotide tokenisation and base-resolution predictions over sequences of up to 1 Mb. The models vary considerably in tokenisation strategy, positional encoding, context length, and parameter count. DNABERT-S further extends DNABERT-2 through curriculum contrastive fine-tuning to produce species-aware embeddings. Key model characteristics are summarised in Table 2. NTv1 was retained only for pooling, timing, and dimensionality reduction experiments because its 1,000 -token context window limited its applicability in robustness and long-sequence analyses and overlapped substantially with the newer NTv2 models.

**Table 2.**
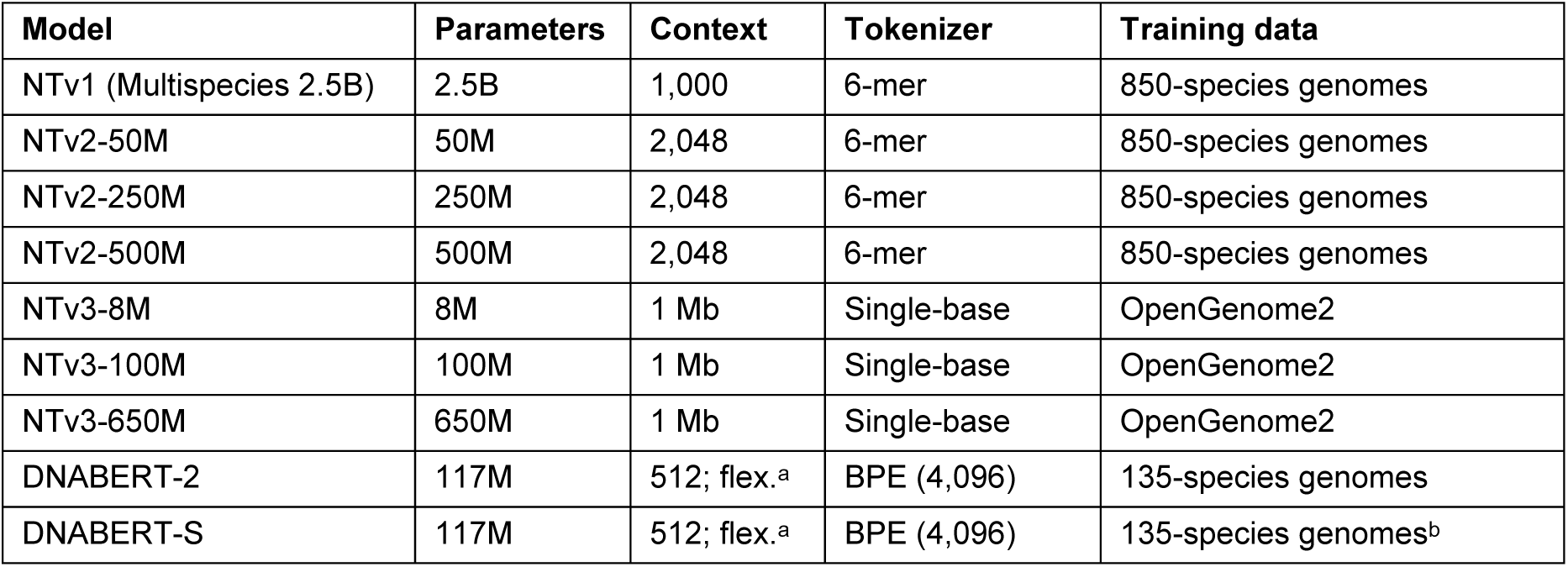
Overview of the evaluated models. Context length is given in tokens unless stated otherwise. ᵃALiBi positional encoding removes the hard input-length constraint; effective context is limited by GPU memory. ᵇDNABERT-S is contrastively fine-tuned from DNABERT-2; architecture and tokenizer are identical.

The robustness and long-sequence experiments were run on a representative subset of four models, namely DNABERT-2, NTv2-50M, NTv2-250M, and NTv3-650M. The subset spans all three tokenisation strategies evaluated here, that is byte-pair, k-mer, and single-nucleotide, and within each family it takes the variant that performed best in the pooling comparison (Section 3.1). DNABERT-S was omitted because it is architecturally identical to DNABERT-2, and the two NTv2 sizes were both retained deliberately, to test whether parameter count alone changes behaviour under corruption and on long genomes.

### 2.4 Embedding generation and pooling

All models were loaded from their official Hugging Face repositories without quantisation or parameter efficient fine-tuning, transferred to GPU, and set to evaluation mode. The study was restricted to forward-pass embedding extraction with frozen weights, so no model parameter was updated at any point. For DNABERT-2 and DNABERT-S, a Triton-syntax incompatibility in their Flash Attention implementation was resolved by replacing the deprecated trans_b argument with an explicit transpose, using publicly available re-published checkpoints in which this fix is already applied. Sequences were loaded from FASTA and sorted by length before being grouped into batches. Every sequence in a batch is padded to the length of the longest member, so batching sequences of very different lengths wastes computation on padding. Sorting places sequences of similar length together and keeps that waste small. Padded positions are excluded by the attention mask and take no part in pooling, so this affects run time only and not the embeddings.

Tokenisation was performed in batches. For all models except NTv3, sequences within a batch were padded to the longest member (padding=’longest’). NTv3 additionally required structural special tokens to be disabled (add_special_tokens=False) and padding to a multiple of 128 (pad_to_multiple_of=128), consistent with its convolutional downsampling architecture. Handling of ambiguous bases differed by family. DNABERT-2 and DNABERT-S mapped N to the [UNK] token, whereas the NT models used an explicit N token.

Each forward pass produced a per-token hidden-state tensor H ∈ R^(B×L×d), where B is batch size, L the padded token length, and d the hidden dimension. This was reduced to a single fixed-length embedding per sequence by one of five pooling configurations. Mean pooling and max pooling were each applied twice, once over all tokens and once with the structural special tokens excluded, and the fifth configuration used the [CLS] token alone. Special-token exclusion was implemented by zeroing the corresponding positions in the attention mask before pooling. For NTv1 and NTv2 this removed [CLS] only, because these models use no [SEP] token, whereas for DNABERT-2 and DNABERT-S both [CLS] and [SEP] were removed.

The three NTv3 models were a special case. Because they were run without structural special tokens (add_special_tokens=False), they had no [CLS] or [SEP] position, so neither the special-token-excluded variants nor CLS pooling was defined. NTv3 was therefore evaluated with mean and max pooling over all tokens only. This gave five configurations for the six NTv1, NTv2, and DNABERT models and two for the three NTv3 models, that is 36 model and pooling combinations in total, and the configurations that do not apply are marked as not applicable in Table 3. Embeddings were serialised to disk for subsequent indexing.

**Table 3.** Accuracy for all models across pooling configurations on the 33-class dataset (maximum length 2,046 bp). Best per model in bold, overall best marked *. Special tokens are the [CLS] and [SEP] positions, so the excluded-token variants and CLS pooling apply only to models that use them. The three NTv3 models are run without structural special tokens and are marked — for those three configurations.

| Pooling configuration | NTv1 | NTv2-50M | NTv2-250M | NTv2-500M | NTv3-8M | NTv3-100M | NTv3-650M | DNABERT-2 | DNABERT-S |
| --- | --- | --- | --- | --- | --- | --- | --- | --- | --- |
| Mean, all tokens | **0.9109** | **0.9133** | **0.9103** | **0.9063** | 0.6509 | **0.9174** | 0.6745 | **0.8855** | **0.8735** |
| Mean, special tokens excluded | 0.9097 | 0.9127 | 0.9099 | 0.9059 | — | — | — | 0.8848 | 0.8733 |
| Max, all tokens | 0.9030 | 0.8673 | 0.8343 | 0.8701 | **0.8210** | 0.8271 | **\*0.9265** | 0.7198 | 0.7907 |
| Max, special tokens excluded | 0.9028 | 0.8667 | 0.8331 | 0.8731 | — | — | — | 0.7495 | 0.8000 |
| CLS token only | 0.8655 | 0.7584 | 0.7651 | 0.7451 | — | — | — | 0.7053 | 0.7713 |

### 2.5 Long-sequence handling

For sequences exceeding a model’s context window, an overlapping token chunking strategy split the input into chunks of 2,046 tokens with an overlap of 250 tokens between consecutive chunks, re-adding the appropriate special tokens per chunk according to model type. DNABERT-2 used a smaller chunk size of 1,200 tokens, reflecting its byte-pair tokenisation, with the same overlap. Chunk embeddings were pooled to a sequence-level representation. The overlap covers about 12% of each chunk, and tokens inside an overlap region contribute to two chunks, so they are counted twice under chunk-level pooling and boundary regions carry correspondingly more weight than the rest of the sequence. This is one reason why chunk-level and sequence-level scoring are not equivalent, a point returned to in Section 4. NTv3-650M was exempt from chunking, as its context window (up to 1 Mb on an A100 GPU) accommodated all sequences in the long-sequence dataset without subdivision.

### 2.6 Dimensionality reduction

To assess whether the full embedding dimension is necessary for classification, principal component analysis (PCA) was fitted on the training embeddings and applied to both training and test embeddings. The number of retained components was set as a fraction of the original dimension, from the full dimension (ratio 1.00) down to 2.5%. A new nearest-neighbour (NN) index was rebuilt at each level and accuracy re-evaluated. This experiment used the 33-class dataset and each model’s best pooling strategy.

### 2.7 Nearest-neighbour classification

Classification was performed by NN search with FAISS [19]. A separate index was built from the training embeddings for each model and pooling combination. Embeddings were L2-normalised (faiss.normalize_L2) prior to indexing, enabling inner-product search (IndexFlatIP) as a proxy for cosine similarity. For each test sequence, the single nearest training neighbour was retrieved (k = 1) and its taxonomic label assigned as the prediction. Because the index is built from the labelled training sequences, this protocol is an NN probe on frozen representations, not pure zero-shot classification in the strict sense, in which no labelled example of a class is available. BLAST received exactly the same labelled sequences in its database, so the two methods were given identical supervision and differed only in how a query was compared with that reference set. In retrieval terms the training split is the indexed reference collection and the test split provides the queries, so “training” here denotes membership of that collection and no parameter update.

### 2.8 BLAST baseline

BLAST was installed locally and a custom nucleotide database was built from the training FASTA, ensuring a controlled and fair comparison with the embedding-based approaches. Two task settings were evaluated, i.e. blastn (optimised for more distant comparisons) and megablast (optimised for highly similar sequences, and substantially faster). The effect of the max_target_seqs parameter (1, 50, 100) on accuracy and runtime was assessed. All main results used max_target_seqs = 1. The taxonomic label of the top hit was taken as the prediction.

### 2.9 Evaluation metrics

Accuracy was the primary metric, complemented by macro-averaged F1-score, precision, and recall. For the long-sequence dataset, balanced accuracy and the Matthews correlation coefficient (MCC) were additionally reported, because the chunking strategy produces an unequal number of chunks per sequence and hence an imbalanced test distribution. All metrics were point estimates from a single fixed split, so accuracies were accompanied by 95% Wilson binomial confidence intervals. With 4,950 test sequences in the refined primary dataset, the interval half-width near an accuracy of 0.93 was about 0.007, so differences below roughly 0.01 fell within sampling error and were not interpreted as meaningful. On the long-sequence dataset, which comprises 256 test sequences, the corresponding half-width was about 0.026.

### 2.10 Compute environment

Experiments were run in Google Colab. Final setup used an NVIDIA A100 GPU. Key software used: Python 3.12.13, Transformers 4.57.3, Hugging Face Hub 0.36.2, Triton 3.6.0, FAISS-CPU 1.13.2, einops 0.8.2, Biopython 1.87, and BLAST 2.12.0.

## 3. Results

### 3.1 Pooling strategy and model comparison

Pooling configurations were compared on the primary dataset (maximum length 2,046 bp), five for the NTv1, NTv2, and DNABERT models and two for the NTv3 models. Table 3 and Figure 1 report the corresponding accuracies on the refined 33-class dataset. Mean pooling over all tokens was the strongest configuration for the NTv1, NTv2, and DNABERT models and also for NTv3-100M, whereas max pooling was best for NTv3-8M and NTv3-650M. Among the six models that have a [CLS] token, CLS pooling was the weakest configuration for every one of them. Excluding the special tokens before mean pooling changed accuracy negligibly, indicating that these tokens contribute little to the mean-pooled representation in the absence of fine-tuning. NTv3-650M achieved the highest overall accuracy (0.93) under max pooling. The two NTv3 models that favour max pooling also show by far the largest spread between configurations. NTv3-650M falls from 0.93 under max pooling to 0.67 under mean pooling, and NTv3-8M from 0.82 to 0.65, so for single-nucleotide tokenisation the choice of pooling matters more than the choice of model.

**Figure 1.**
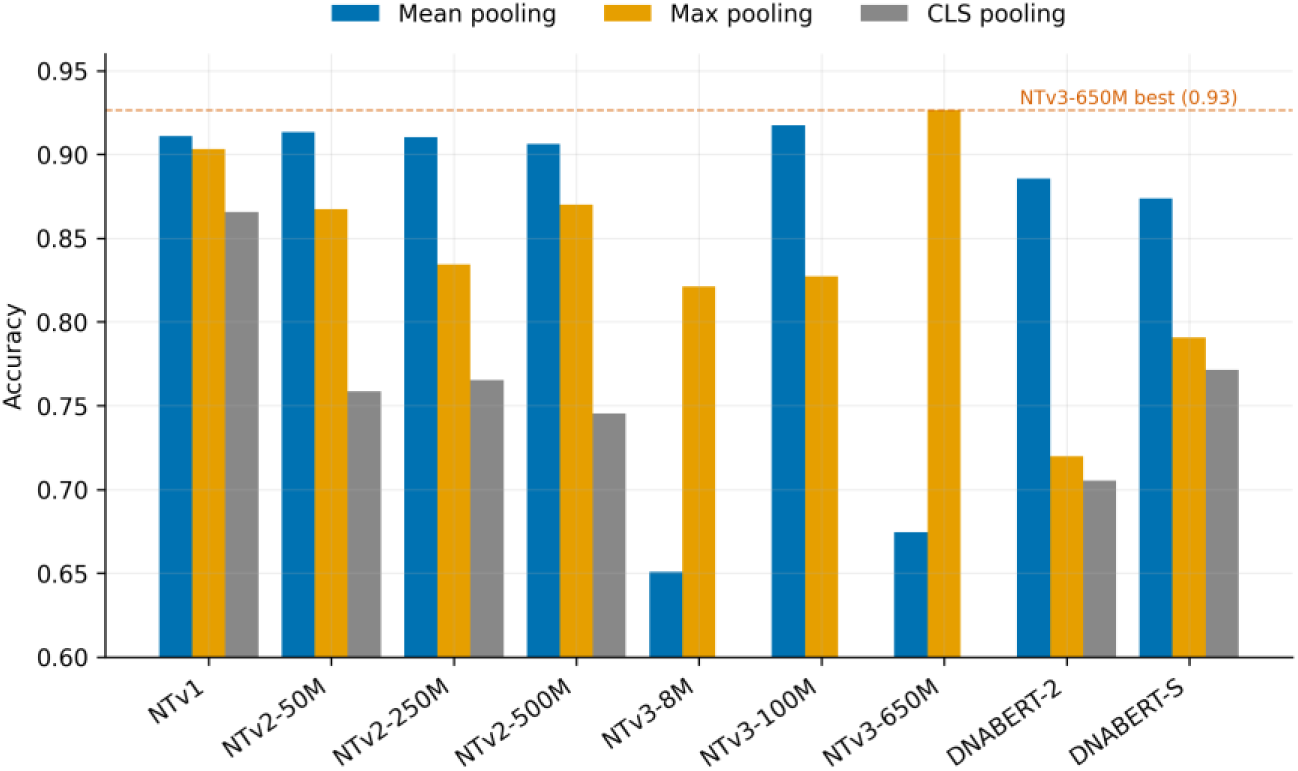
Classification accuracy by pooling strategy (mean, max, and CLS over all tokens) for all nine models on the refined 33-class dataset. The dashed line marks the best result (NTv3-650M, max pooling, 0.93). No CLS bar is shown for the three NTv3 models, which have no [CLS] token.

### 3.2 Taxonomic ambiguity and dataset refinement

Systematic inspection of per-model confusion matrices revealed several class groups that were consistently misclassified across all nine models as well as BLAST. The dominant patterns involved parent and child taxonomic relationships and overlapping designations. Quoted below are the RVDB label strings themselves, which is why the same virus appears under more than one name, namely, Porcine reproductive and respiratory syndrome virus / PRRSV-2 / Betaarterivirus americense; Human immunodeficiency virus / HIV-1; Hepacivirus hominis and its Hepatitis C subtypes; Enterovirus B and its member species (Coxsackievirus B5, Echovirus E30, and others); and human respiratory syncytial virus / Human respiratory syncytial virus A. The most frequent misclassification pairs are summarised in Table 4 and Figure 2. Removing seven such ambiguous classes reduced the dataset from 40 to 33 classes and substantially improved accuracy for all methods, including BLAST. These seven labels were afterwards reconciled against the International Committee on Taxonomy of Viruses (ICTV), which confirmed that they corresponded to five species and identified one further redundant pair that the 33-class dataset still contained (Section 4, Table 11).

**Figure 2.**
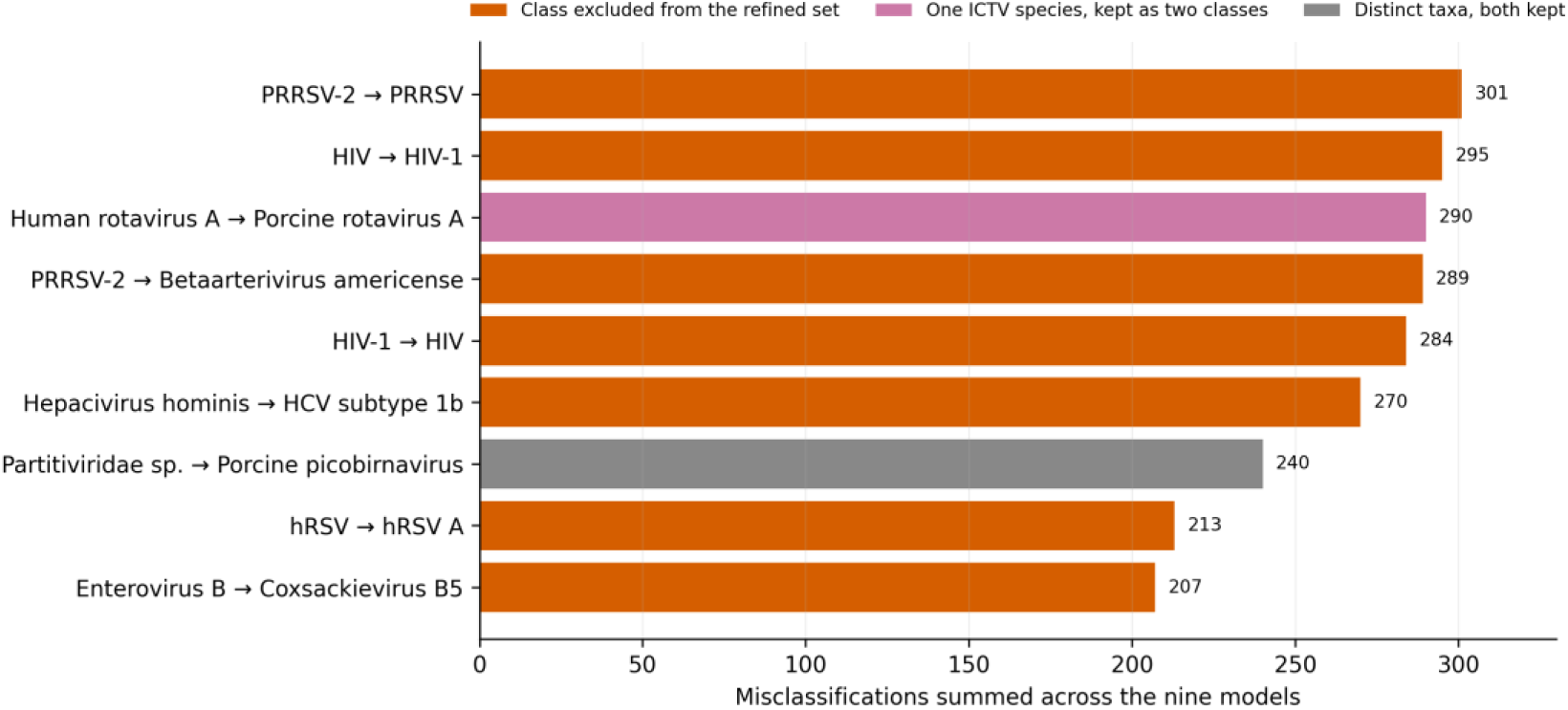
Most frequent misclassifications on the 40-class dataset. Horizontal bars give the number of misclassifications summed across the nine models for each true-to-predicted class pair. Red marks pairs involving a class excluded from the refined 33-class set. Purple marks the rotavirus pair, which was kept as two classes although both labels correspond to a single ICTV species. Grey marks the one pair in which both classes are distinct taxa (Table 11).

**Table 4.** Selected misclassification pairs, sorted by total occurrences across the nine models. “Models” gives the number of models (of 9) in which the mismatch appeared. Excluded classes are shown in bold. The Human rotavirus A and Porcine rotavirus A labels were kept as separate classes although both correspond to a single ICTV species (Table 11).

| True class | Predicted class | Total | Models |
| --- | --- | --- | --- |
| **PRRSV-2** | **PRRSV** | 301 | 9/9 |
| **Human immunodeficiency virus** | HIV-1 | 295 | 9/9 |
| Human rotavirus A | Porcine rotavirus A | 290 | 9/9 |
| **PRRSV-2** | **Betaarterivirus americense** | 289 | 9/9 |
| HIV-1 | **Human immunodeficiency virus** | 284 | 9/9 |
| **Hepacivirus hominis** | Hepatitis C virus subtype 1b | 270 | 9/9 |
| Partitiviridae sp. | Porcine picobirnavirus | 240 | 9/9 |
| **human respiratory syncytial virus** | Human respiratory syncytial virus A | 213 | 9/9 |
| **Enterovirus B** | Coxsackievirus B5 | 207 | 9/9 |

### 3.3 Effect of model scale

Larger models did not consistently outperform smaller ones (Table 3). On the 33-class dataset, NTv1 (2.5B), NTv2-50M, NTv2-250M, and NTv2-500M all clustered near 0.91, with a spread of about 0.007 that lies within the sampling error of this test set. Within the NTv3 family, the pattern differed. NTv3-650M was best overall (0.93, max pooling), NTv3-100M second (0.92, mean pooling), and NTv3-8M markedly weaker (0.82 max, 0.65 mean). Both DNABERT models were the weakest overall, and the DNABERT-S (0.87) did not surpass DNABERT-2 (0.89), indicating that species-aware model does not transfer to an advantage on this viral identification task.

### 3.4 Processing time and batch size

Embedding generation time was measured across batch sizes of 4 to 1,000 on the primary test set (maximum 2,046 bp; mean pooling). Some large batches were infeasible for the largest models due to A100 memory limits. NTv3-8M was fastest overall, reaching 6.45 s at batch size 500, while NTv1 was slowest (best 242.33 s at batch size 100), reflecting its large parameter count and 2,560-dimensional embeddings. Among NTv2 models, NTv2-50M offered the best trade-off between speed and accuracy (17.82 s at batch size 100). DNABERT-2 stabilised at about 22 to 24 s versus about 60 to 71 s for NTv2-500M at larger batches. Based on these measurements, batch size 100 was adopted for NTv1/NTv2 and 200 for NTv3/DNABERT (Table 5, Figure 3).

**Figure 3.**
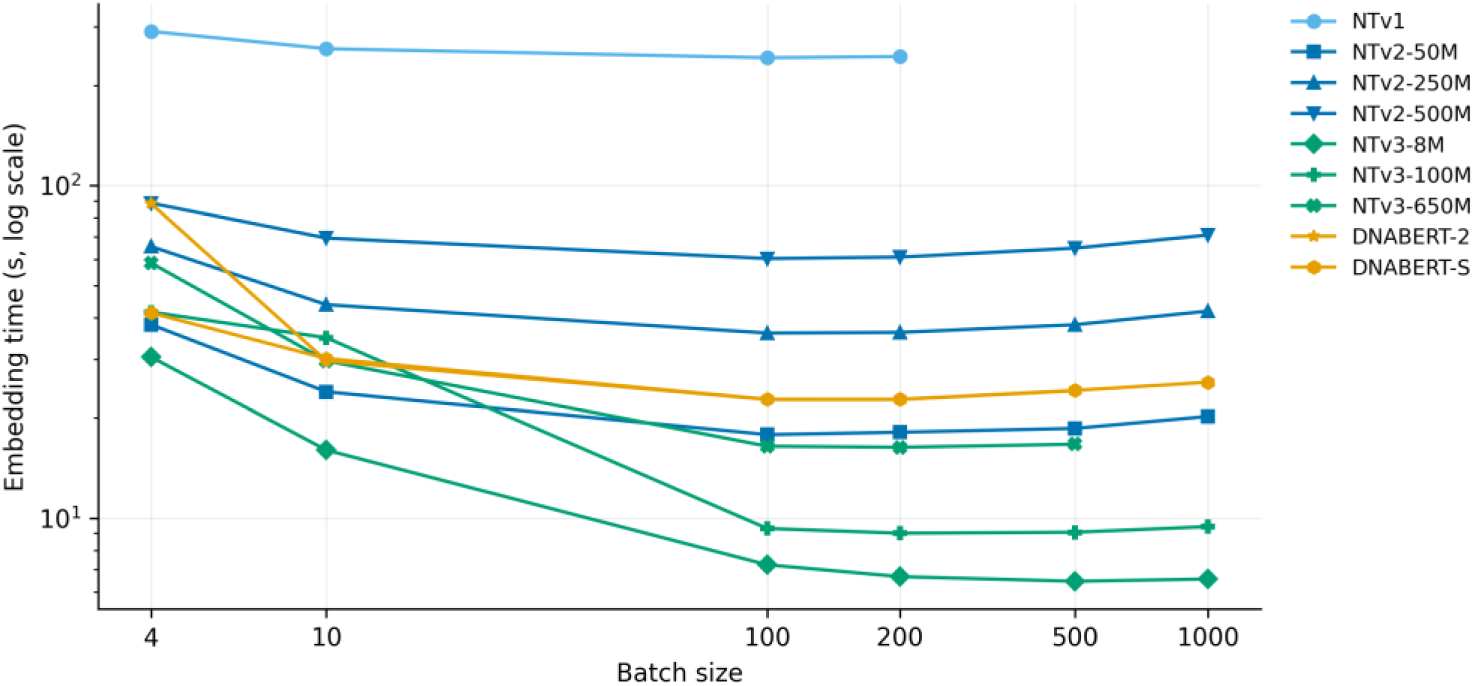
Embedding generation time versus batch size (primary test set, maximum 2,046 bp; mean pooling; NVIDIA A100). Log-log axes; missing points indicate out-of-memory configurations.

**Table 5.** Embedding-generation time (s) as a function of batch size (primary test set, maximum 2,046 bp; mean pooling). “—” indicates an out-of-memory error on the A100.

| Model | 4 | 10 | 100 | 200 | 500 | 1000 |
| --- | --- | --- | --- | --- | --- | --- |
| NTv1 | 290.79 | 257.96 | 242.33 | 244.35 | — | — |
| NTv2-50M | 38.16 | 23.96 | 17.82 | 18.12 | 18.61 | 20.21 |
| NTv2-250M | 65.57 | 43.81 | 35.99 | 36.15 | 38.13 | 41.91 |
| NTv2-500M | 88.48 | 69.49 | 60.32 | 60.93 | 64.84 | 70.97 |
| NTv3-8M | 30.48 | 16.01 | 7.23 | 6.66 | 6.45 | 6.55 |
| NTv3-100M | 41.63 | 34.83 | 9.30 | 9.00 | 9.06 | 9.42 |
| NTv3-650M | 58.48 | 29.68 | 16.45 | 16.33 | 16.68 | — |
| DNABERT-2 | 88.22 | 29.68 | 22.71 | 22.71 | 24.12 | — |
| DNABERT-S | 41.38 | 30.17 | 22.74 | 22.73 | 24.19 | 25.59 |

### 3.5 Dimensionality reduction

PCA showed that the embeddings were highly compressible (Table 6, Figure 4). For most models, retaining between 80 and 90% of the original dimensions left accuracy unchanged or marginally higher than the full-dimension baseline. These small gains were within sampling error. For NTv3-650M, for example, the apparent gain at 80% retention corresponded to three of the 4,950 test sequences. NTv2-50M was the one model with a larger change, rising from 0.91 (512 dims) to 0.92 at 60% (307 dims), a difference of about 30 sequences. NTv3-650M retained about 0.92 accuracy even at 25% of its original dimension (384 dims), a fourfold compression of the 1,536-dimensional space at negligible accuracy cost, which is relevant to storage and search efficiency at scale. Across all models, accuracy at 25% remained well above chance, and for most NT models above 0.90.

**Figure 4.**
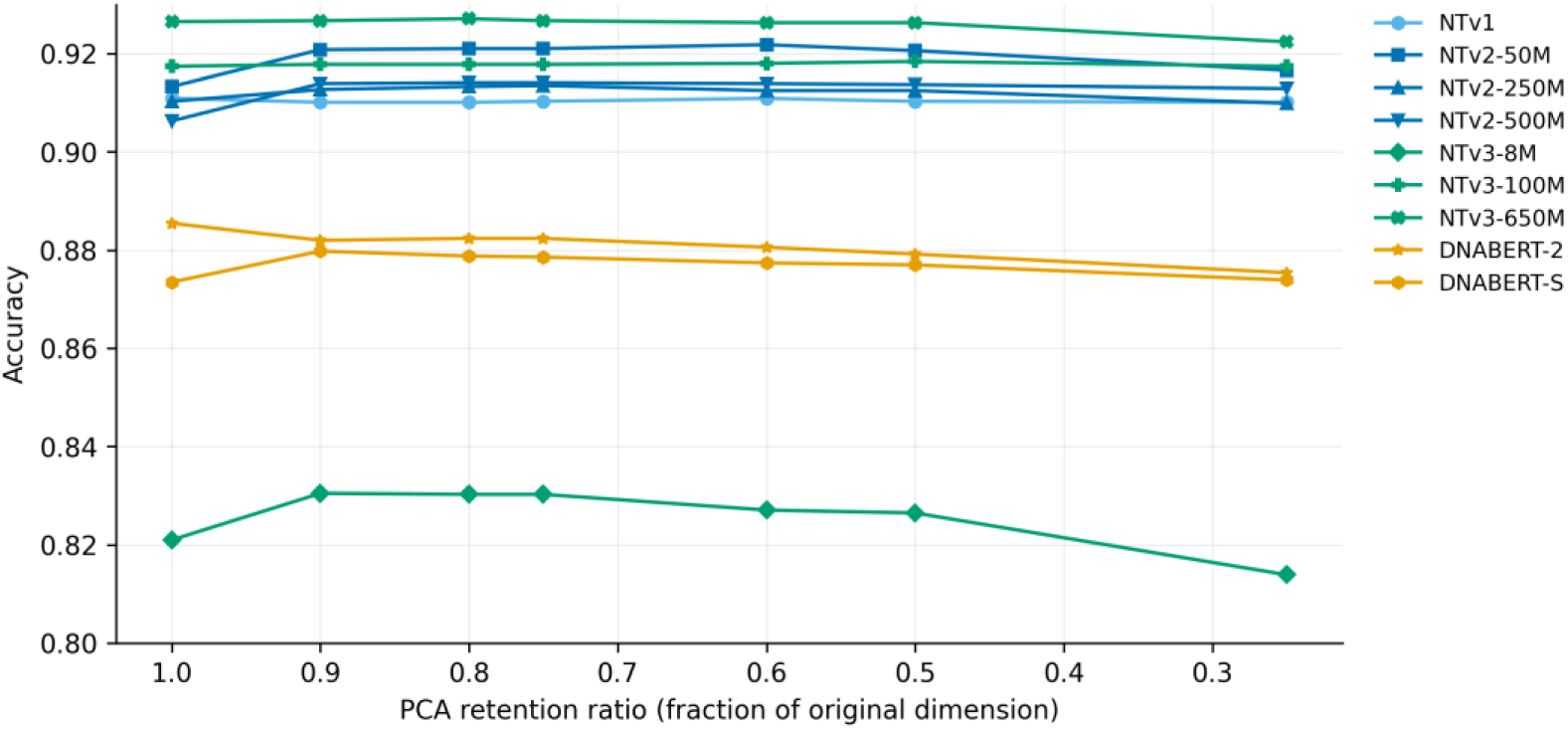
Classification accuracy versus PCA retention ratio on the refined 33-class dataset.

**Table 6.** Accuracy after PCA-based dimensionality reduction on the 33-class dataset. “Orig. dim” is the original embedding dimension; remaining columns give accuracy at the indicated retention ratios.

| Model | Orig. dim | 1.00 | 0.90 | 0.80 | 0.75 | 0.60 | 0.50 | 0.25 |
| --- | --- | --- | --- | --- | --- | --- | --- | --- |
| NTv1 | 2560 | 0.9109 | 0.9101 | 0.9101 | 0.9103 | 0.9109 | 0.9103 | 0.9101 |
| NTv2-50M | 512 | 0.9133 | 0.9208 | 0.9210 | 0.9210 | 0.9218 | 0.9206 | 0.9166 |
| NTv2-250M | 768 | 0.9103 | 0.9127 | 0.9133 | 0.9135 | 0.9125 | 0.9125 | 0.9099 |
| NTv2-500M | 1024 | 0.9063 | 0.9139 | 0.9141 | 0.9141 | 0.9139 | 0.9137 | 0.9129 |
| NTv3-8M | 256 | 0.8210 | 0.8305 | 0.8303 | 0.8303 | 0.8271 | 0.8265 | 0.8139 |
| NTv3-100M | 768 | 0.9174 | 0.9178 | 0.9178 | 0.9178 | 0.9180 | 0.9184 | 0.9174 |
| NTv3-650M | 1536 | 0.9265 | 0.9267 | 0.9271 | 0.9267 | 0.9263 | 0.9263 | 0.9224 |
| DNABERT-2 | 768 | 0.8855 | 0.8820 | 0.8824 | 0.8824 | 0.8806 | 0.8792 | 0.8754 |
| DNABERT-S | 768 | 0.8735 | 0.8798 | 0.8788 | 0.8786 | 0.8774 | 0.8770 | 0.8739 |

### 3.6 Comparison with BLAST

Varying max_target_seqs across 1, 50, and 100 had no meaningful effect on BLAST accuracy, but it increased runtime severalfold. For blastn on the 40-class dataset, runtime rose from 166.75 s with a single target sequence to 690.67 s with one hundred. All main results therefore used max_target_seqs = 1. The comparison against the best GFM configuration (NTv3-650M, max pooling) is given in Table 7 and Figure 5. BLAST (blastn) achieved the highest accuracy on both datasets (0.91 on the 40-class and 0.97 on the 33-class dataset), outperforming NTv3-650M (0.88 and 0.93) and megablast (0.85 and 0.90). On the 33-class dataset the 95% confidence intervals do not overlap (blastn 0.962 to 0.972, NTv3-650M 0.919 to 0.933, megablast 0.891 to 0.907). megablast reached the highest precision but the lowest recall, reflecting its conservative matching for near-identical sequences, and ran roughly three to four times faster than blastn. On the 33-class dataset, NTv3-650M closed the gap to blastn to 0.04 accuracy and clearly surpassed megablast.

**Figure 5.**
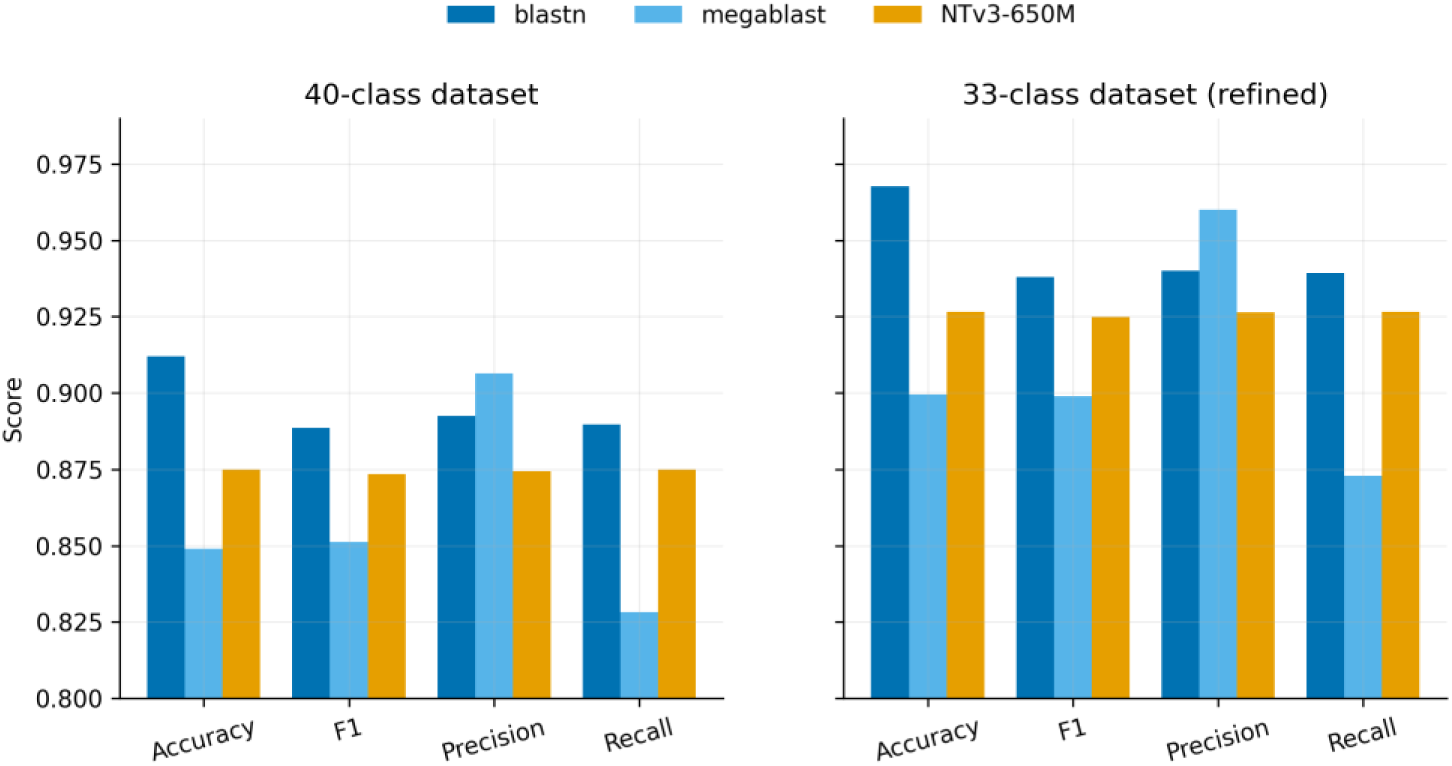
BLAST (blastn, megablast) versus the best foundation model (NTv3-650M, max pooling) on the 40-class and refined 33-class datasets across four metrics.

**Table 7.** Comparison of BLAST with the best GFM configuration (NTv3-650M, max pooling) on the 40- and 33-class datasets (max_target_seqs = 1).

| Dataset | Method | Accuracy | F1 | Precision | Recall |
| --- | --- | --- | --- | --- | --- |
| 40 classes | blastn | **0.9120** | 0.8886 | 0.8925 | 0.8898 |
| 40 classes | megablast | 0.8490 | 0.8512 | 0.9064 | 0.8283 |
| 40 classes | NTv3-650M (max) | 0.8750 | 0.8735 | 0.8744 | 0.8750 |
| 33 classes | blastn | **0.9677** | 0.9379 | 0.9401 | 0.9392 |
| 33 classes | megablast | 0.8994 | 0.8989 | 0.9600 | 0.8729 |
| 33 classes | NTv3-650M (max) | 0.9265 | 0.9249 | 0.9263 | 0.9265 |

### 3.7 Robustness to mutation and masking

Robustness was assessed on the 6 × 6 grid of mutation and masking rates for BLAST (blastn) and the four representative models defined in Section 2.3 (DNABERT-2, NTv2-50M, NTv2-250M, NTv3-650M). BLAST was strongly robust. Accuracy remained above 0.96 up to 5% mutation without masking, 0.94 at 20% mutation without masking, and 0.93 at 20% masking without mutation. Only the combination of high mutation and high masking degraded substantially (0.56 at 20% mutation with 20% masking).

In contrast, all foundation models were far more sensitive to masking than mutation (Figure 6), the practical opposite of BLAST. The effect was severe for the k-mer and BPE models. NTv2-50M fell from 0.91 to 0.68 at 1% masking and to 0.14 at 5%, and DNABERT-2 fell to 0.76 at 1% masking. NTv3-650M was the most masking-robust foundation model, retaining 0.91 at 2% masking and 0.85 at 5% masking (0% mutation), but it too collapsed at higher masking (0.59 at 10%, 0.21 at 20%) and dropped below 0.90 at 2% mutation (0.89). Its greater robustness is plausibly attributable to single-nucleotide tokenisation, under which an N replaces a single token rather than corrupting an entire k-mer or subword unit, limiting the propagation of the masking effect. Full accuracy grids are given in Tables 8 and 9 (the remaining models follow the same masking-dominated pattern).

**Figure 6.**
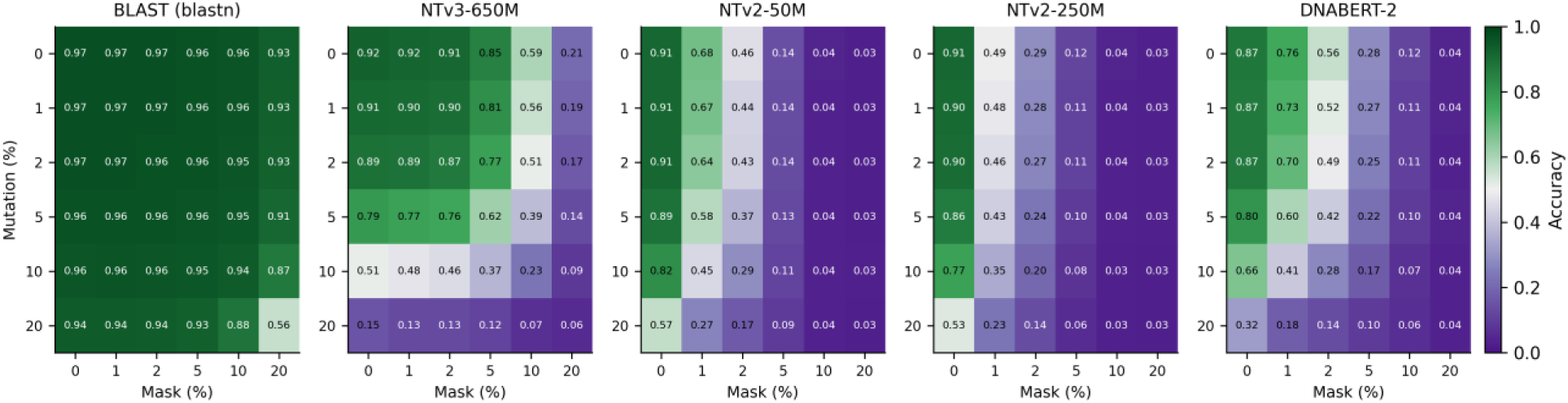
Robustness to mutation and masking on the refined 33-class dataset. Accuracy over the 6×6 grid of mutation (rows) and masking (columns) rates for BLAST and four representative models, shared colour scale.

**Table 8.** BLAST (blastn) accuracy on the 33-class dataset. Rows: mutation rate (%); columns: masking rate (%).

| Mut Mask | 0 | 1 | 2 | 5 | 10 | 20 |
| --- | --- | --- | --- | --- | --- | --- |
| 0 | 0.9677 | 0.9671 | 0.9667 | 0.9644 | 0.9572 | 0.9313 |
| 1 | 0.9663 | 0.9675 | 0.9667 | 0.9626 | 0.9574 | 0.9285 |
| 2 | 0.9661 | 0.9659 | 0.9646 | 0.9632 | 0.9549 | 0.9251 |
| 5 | 0.9640 | 0.9646 | 0.9632 | 0.9576 | 0.9515 | 0.9127 |
| 10 | 0.9570 | 0.9598 | 0.9552 | 0.9525 | 0.9382 | 0.8703 |
| 20 | 0.9398 | 0.9384 | 0.9358 | 0.9275 | 0.8806 | 0.5608 |

**Table 9.** NTv3-650M accuracy on the 33-class dataset. Rows: mutation rate (%); columns: masking rate (%).

| Mut Mask | 0 | 1 | 2 | 5 | 10 | 20 |
| --- | --- | --- | --- | --- | --- | --- |
| 0 | 0.9244 | 0.9170 | 0.9131 | 0.8495 | 0.5945 | 0.2077 |
| 1 | 0.9073 | 0.9024 | 0.8980 | 0.8139 | 0.5556 | 0.1941 |
| 2 | 0.8899 | 0.8851 | 0.8695 | 0.7745 | 0.5105 | 0.1693 |
| 5 | 0.7907 | 0.7707 | 0.7580 | 0.6160 | 0.3881 | 0.1362 |
| 10 | 0.5083 | 0.4760 | 0.4570 | 0.3723 | 0.2307 | 0.0879 |
| 20 | 0.1533 | 0.1311 | 0.1299 | 0.1158 | 0.0697 | 0.0576 |

### 3.8 Long sequences

On the long-sequence dataset (84,587 to 409,110 bp, about 4 training and about 16 test sequences per class, 256 test sequences in total), NTv3-650M was the strongest foundation model (accuracy 0.95, balanced accuracy 0.95, MCC 0.95), ahead of NTv2-50M (0.85), NTv2-250M (0.83), and DNABERT-2 (0.79). NTv3-650M and BLAST processed each sequence whole, whereas the three shorter-context models were scored over chunks, so part of the difference may reflect the unit of evaluation (Section 4). BLAST (blastn) achieved the highest accuracy overall (0.98). The gap to NTv3-650M was about 0.02, which corresponds to six of the 256 test sequences, and the 95% confidence intervals overlap (NTv3-650M 0.920 to 0.973, blastn 0.950 to 0.989). On a test set of this size, the narrowing relative to the primary dataset is therefore indicative, and confirming it would require a paired test on per-sequence outcomes and a larger long-sequence collection (Figure 7, Table 10). NTv3-650M processed each genome without chunking thanks to its 1 Mb context window.

**Figure 7.**
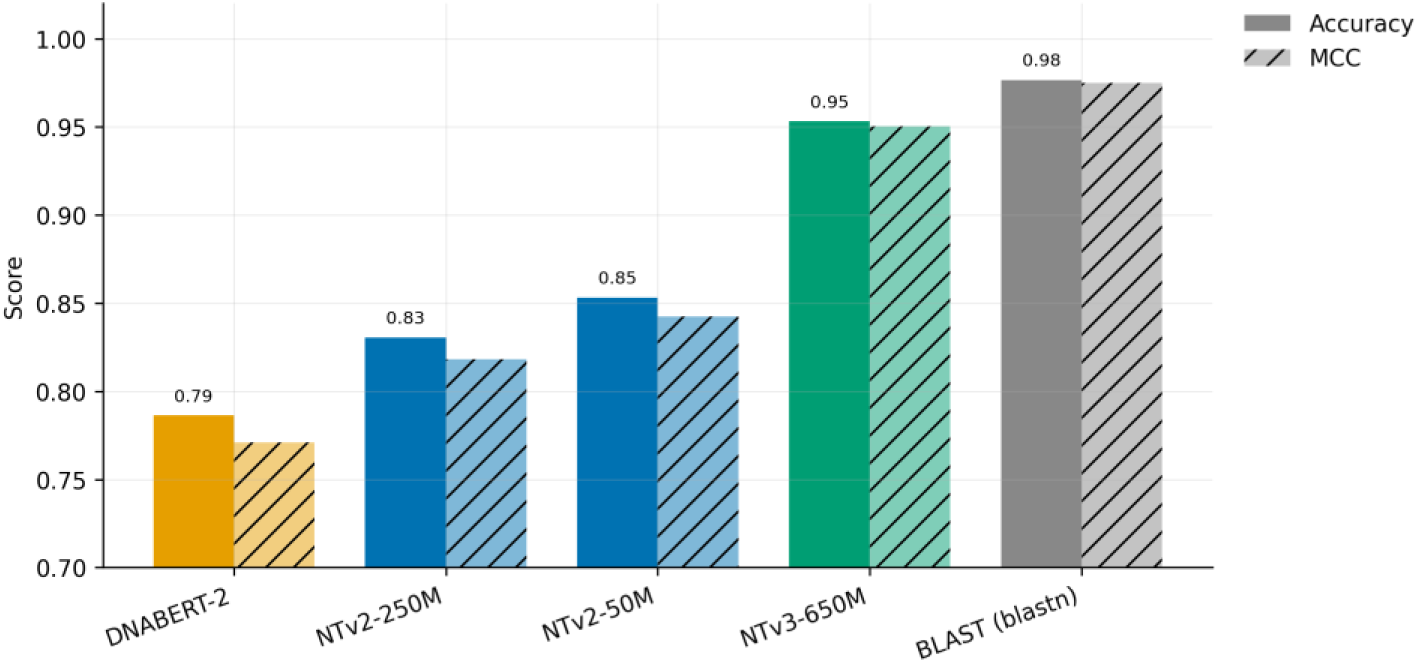
Long-sequence dataset (84.6 to 409 kb; 16 classes). Accuracy and Matthews correlation coefficient for four foundation models and BLAST.

**Table 10.** Classification results on the long-sequence dataset (16 classes; about 4 training and about 16 test sequences per class; 256 test sequences). ᵃEvaluated over overlapping chunks rather than whole sequences, because the model context window is shorter than these genomes; metrics for these rows are therefore not computed on the same unit as for NTv3-650M and blastn.

| Model | Accuracy | Bal. Acc. | MCC | F1 | Precision | Recall |
| --- | --- | --- | --- | --- | --- | --- |
| DNABERT-2<sup>a</sup> | 0.7864 | 0.8170 | 0.7711 | 0.8127 | 0.8139 | 0.8170 |
| NTv2-50M<sup>a</sup> | 0.8532 | 0.8699 | 0.8423 | 0.8689 | 0.8705 | 0.8699 |
| NTv2-250M<sup>a</sup> | 0.8306 | 0.8443 | 0.8181 | 0.8423 | 0.8435 | 0.8443 |
| NTv3-650M | 0.9531 | 0.9531 | 0.9504 | 0.9534 | 0.9583 | 0.9531 |
| BLAST (blastn) | **0.9766** | 0.9766 | 0.9751 | 0.9765 | 0.9776 | 0.9766 |

**Table 11.**
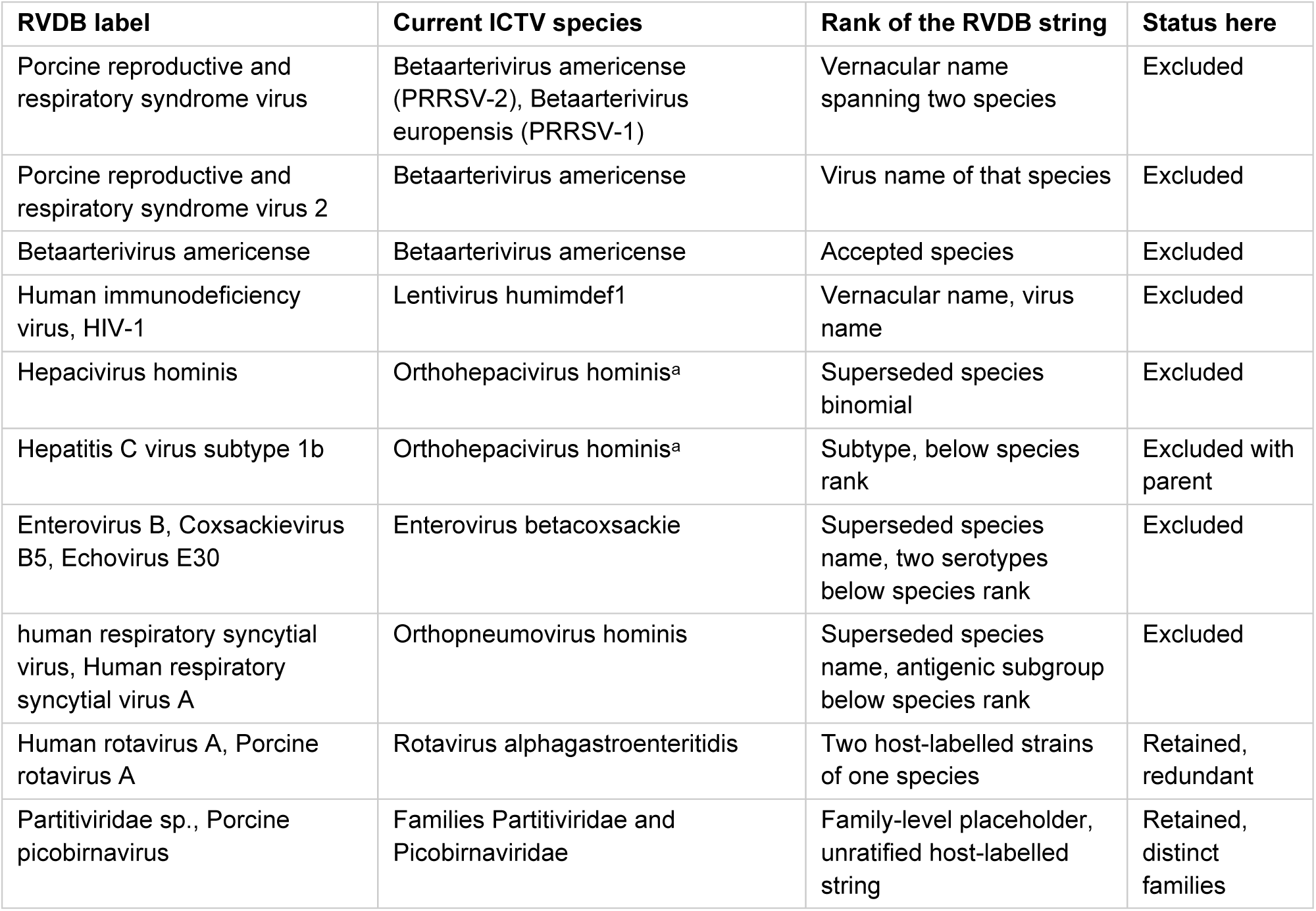
Reconciliation of the RVDB labels involved in the most frequent confusions against the ICTV taxonomy (MSL41). Rank refers to the status of the RVDB string itself, not to the virus it denotes. ᵃA reorganisation of the Flaviviridae that would further adjust genus assignments in this group has been proposed but is not yet ratified, and some ICTV Report chapters still list the earlier name Hepacivirus hominis.

## 4. Discussion

This study set out to characterise how GFMs compare with alignment-based search for viral sequence identification, using a controlled, RVDB-derived protocol in which frozen embeddings were classified by NN retrieval. Three findings anchor the discussion. Alignment (BLAST) remains the stronger method on a closed reference database, yet the best foundation model (NTv3-650M) comes within a small margin without any task-specific training. The embedding space is highly compressible and the accuracy gap narrows on long genomes. All foundation models remain markedly less robust to base-level corruption than BLAST.

### Overall performance

On the 33-class dataset, blastn reached 0.97 against 0.93 for the best foundation model, NTv3-650M. The two pipelines received the same supervision. Both indexed the same 350 training sequences per class, and both can only return a label present in that reference set, so neither is open-set at prediction time.

### What differs is how a query is compared with the reference

BLAST compares nucleotides and scores local matches, which is close to the ideal operation for retrieval from a curated database of near-relatives. The foundation model has to route the same comparison through a representation that was fixed in advance by generic pre-training and never adapted to this database, and it must separate 33 classes within that space. The broad prior that makes foundation models attractive for novel sequences grants no advantage on this comparison, because the model carries far more information than the task requires while still having to resolve fine distinctions among closely related references. Reaching within about 0.04 accuracy of an aligner from fixed, task-agnostic representations, with no parameter updates and no task-specific training, is therefore considered as a strong result. The gap was smaller on long sequences (0.95 against 0.98), where the 1 Mb context window of NTv3 let it encode entire genomes without chunking and preserve global context that alignment otherwise captures only piecewise. That attribution is plausible but not isolated, because the two datasets differ in more than sequence length. The long-sequence benchmark indexed about 4 reference sequences per class (instead of 350), and its classes were drawn from far more divergent viruses, such as Monkeypox virus, Variola virus, and African swine fever virus, instead of the closely related subtypes that dominate the primary dataset.

### Pooling and dimensionality

Mean pooling was the best configuration for six of the nine models and CLS pooling was the worst for every model that has a [CLS] token. This matches an independent benchmark of five DNA foundation models on zero-shot embeddings, in which mean token pooling significantly outperformed both CLS and maximum pooling for sequence classification, for the NT in 42 of 52 datasets [21]. The possible explanation is that, without task-specific fine-tuning, the [CLS] token does not accumulate a discriminative sequence-level representation. The same effect is long established for text encoders, where untuned BERT sentence embeddings underperform simple averaging and motivated dedicated sentence-embedding methods [22,23]. Max pooling was preferable for two of the three single-nucleotide NTv3 models, NTv3-8M and NTv3-650M, but not for NTv3-100M, which favoured mean pooling. Optimal aggregation therefore depends on more than tokenisation granularity alone. Special-token removal was immaterial under mean pooling.

PCA showed the embeddings to be highly compressible, with accuracy retained around 80 to 90% retention and, for NTv3-650M, more than 0.92 accuracy still reached 25% of the original dimension. This compressibility is practically important. A 384-dimensional NTv3-650M embedding enables faster search and lower memory, which might matter for deployment against the full C-RVDB.

### Robustness

The most consequential limitation for real-world use is sensitivity to masking. All foundation models degraded sharply as ambiguous N bases were introduced, whereas BLAST remained above 0.93 even at 20% masking. NTv3-650M was the most robust foundation model, plausibly because single-nucleotide tokenisation limits each N to one token, but still fell below 0.90 by 2% mutation and collapsed at high masking. Because low-quality base calls and ambiguous positions are routine in NGS data, this behaviour, is likely to be the binding constraint on clinical deployment of current frozen embeddings.

### Taxonomic ambiguity

A methodological finding of independent interest is that several RVDB label groups were misclassified identically by every model and by BLAST, which traces those errors to nested and overlapping labels. RVDB inherits organism strings from GenBank, RefSeq, and Third-Party Annotation records without imposing a controlled taxonomy [7], so its labels mix accepted species names, superseded synonyms, dialect virus names, and designations that sit below species rank. We therefore reconciled every label involved in the most frequent confusions against the ICTV taxonomy. Two rules resolve most cases. Species names have been binomial since the 2021 ratification, so several RVDB strings are superseded synonyms of a renamed species [24]. ICTV also assigns no rank below species, which means that serotypes, genotypes, subtypes, and antigenic subgroups are not taxa and cannot serve as mutually exclusive classification targets [25]. Names below follow the current Master Species List, MSL41 [26].

The reconciliation is given in Table 11. It confirms that the seven excluded labels correspond to only five ICTV species. The three PRRSV labels are the single species Betaarterivirus americense. The HIV labels are the single species Lentivirus humimdef1. Hepacivirus hominis and Hepatitis C virus subtype 1b are the species now named Orthohepacivirus hominis together with one of the 67 subtypes recognised within it [27]. Enterovirus B, now Enterovirus betacoxsackie, subsumes the Coxsackievirus B5 and Echovirus E30 serotypes. The respiratory syncytial virus labels are the species Orthopneumovirus hominis together with one of its two antigenic subgroups. The exclusions therefore rest on taxonomic grounds that are independent of our error analysis, which matters because these classes were first identified through the confusion patterns they produced.

The same reconciliation exposes one redundancy that survived into the 33-class dataset. Human rotavirus A and Porcine rotavirus A are host-labelled strains of the single species *Rotavirus alphagastroenteritidis*, whose boundary is set by VP6 sequence identity [28]. This pair produced the third most frequent confusion in Table 4, and because both labels were kept as separate classes, those assignments were taxonomically correct yet scored as errors. Every accuracy reported here, for BLAST as well as for the foundation models, is therefore slightly conservative, and merging the pair is the first change we would make in a follow-up. The other frequent confusion, between Partitiviridae sp. and Porcine picobirnavirus, spans two families and is legitimately kept separate, although neither string is a clean species-level target, because Partitiviridae sp. carries no species assignment and host-named picobirnavirus sequences are known to be phylogenetically heterogeneous. Reconciling labels against ICTV before RVDB annotations are used as supervised targets is thus not an optional refinement but a prerequisite.

### Comparison with prior work

Retrieval over frozen representations has very recently been examined in its own right. Concurrent with this work, Fahmy et al. compared supervised classifiers with FAISS NN retrieval over one-hot, k-mer, FCGR, dna2vec, and DNABERT representations [29]. That study shares our retrieval framing but differs on three axes. Its tasks are genotyping within a single virus (hepatitis C and human papillomavirus) and one binary discrimination task. Furthermore, they used a single first-generation genomic model.

Beyond that, two differences from earlier work are consistent (Table 12). First, to our knowledge no prior GFM study evaluates against RVDB itself. ViraLM and VIRALpre train on RefSeq (supplemented with VirSorter2-curated genomes), and ViroBench draws a curated corpus from RefSeq or GenBank. RVDB is assembled from those same public archives, so the underlying sequences overlap, but its semantic selection, manual curation, and clustering produce a different collection with a different composition of complete genomes, partial CDS, and genomic regions, and that is what a retrieval method is actually indexed against. Second, the dominant task is binary virus versus host detection of metagenomic contigs (ViraLM, VIRALpre), or supervised taxonomy and host classification with a trained head (ViroBench), rather than the multi-class taxonomic retrieval evaluated herein.

**Table 12.** Positioning of this study relative to representative prior work on GFM and non-GFM viral sequence tools. NFM = nucleotide foundation model.

| Study | Reference database | Task | # foundation models | BLAST used as | Base-corruption robustness? | Approach |
| --- | --- | --- | --- | --- | --- | --- |
| This study | C-RVDB v31.0 | Multi-class taxonomic classification (33/40 classes; + long-genome, 16 classes) | 9 | Explicit baseline (blastn, megablast) | Yes (mutation × masking grid) | Frozen-embedding nearest-neighbour retrieval |
| ViraLM [13] | RefSeq + VirSorter2-curated | Binary virus vs. non-virus detection | 1 (DNABERT-2) | Data cleaning only | No | Supervised fine-tuning (2 kb fragments, scores averaged) |
| VIRALpre [14] | RefSeq | Binary virus vs. non-virus detection | 1 (EVO; others compared) | Similarity cut-off test only | No (BLASTn similarity cut-off generalisation) | Fine-tuned; GFM embedding + k-mer fusion |
| NextVir [4] | iCAV (oncoviruses) | Multi-class oncoviral read classification (7 families + human) | 3 (DNABERT-S, NT, HyenaDNA) | Not used | Yes, substitutions and indels (plus contamination) | LoRA fine-tuning + adapter (frozen embeddings also tested) |
| ViroBench [16] | RefSeq / GenBank (curated) | Taxonomy and host prediction; separate generation axis | 66 | Explicit baseline (+ Kraken2) | No (phylogenetic & temporal splits instead) | Supervised classification head |
| Fahmy et al. [29] | Los Alamos HCV database, NCBI | Within-virus genotyping (HCV, HPV) and binary COVID-19 discrimination | 1 (DNABERT; also dna2vec, k-mer, FCGR, one-hot) | Not used | No | FAISS nearest-neighbour retrieval and supervised classifiers |
| Ghorbani et al. [30] | Custom (NGS pipeline) | Virus detection + SNP discovery | 0 | Tuned baseline (E-value, word size) | No | Alignment pipeline with tuned BLAST |
| Wu et al. [31] | RefSeq + real metagenomes | Binary virus detection (tool benchmark) | 0 (CNN/HMM/homology tools) | Homology-only tool included | No | Pre-trained/threshold-tuned tools |

ViraLM and VIRALpre both fine-tune a foundation model end-to-end and use BLASTN only inside their pipelines (ViraLM to clean endogenous viral elements from negatives, VIRALpre as a similarity cut-off to construct a hard test set), whereas we retain BLAST as an explicit baseline. Some alignment-centred virus tools tune BLAST heavily, adjusting E-value thresholds and word size to maximise performance [30]. We did not tune BLAST in this way and varied only max_target_seqs, which had no measurable effect on accuracy. It means the baseline reported here is an untuned configuration that already should provide a strong upper bound.

Prior work that compared foundation models with alignment showed that which method wins depends on the task and on how the data is split. ViroBench evaluated its models under a genus-disjoint split, meaning that no member of a query’s genus is present in the reference data, so no method can succeed simply by retrieving a close relative. Under that split BLAST still predicted the host of a virus well, reaching a macro F1 of 92.50 against 84.56 for the best foundation model, which the authors attributed to similarity-based methods exploiting close matches in the reference database. For taxonomic assignment, the ordering reversed, with BLAST at 47.67 against 75.88 [16]. A plausible reading of that contrast is that host tends to be conserved across related genera, so a match outside the query’s own genus can still carry the correct host label, whereas the same match says much less about where the query belongs in the taxonomy. Taxonomic assignment is the task we evaluate, and the contrast matters for how our own result should be read. Our reference index contains near relatives of every test class, which is the situation alignment handles best, whereas ViroBench removed exactly that advantage and the ordering we report does not survive it. That is a further reason to expect the ordering reported here to change in the open-set setting.

Independent benchmarking of non-GFM tools (DeepVirFinder, VirSorter2, VIBRANT, PPR-Meta, and homology-only methods) similarly finds that learned models can beat homology-only tools but that no single tool dominates and that parameter cut-offs materially change results [31], reinforcing that the comparison target, not just the model, determines the conclusion.

### Methodological considerations

Two aspects of the design warrant particular caution when interpreting the results. First, on the long-sequence dataset the evaluation was not uniform across methods. NTv3-650M and BLAST processed each sequence whole, whereas DNABERT-2, NTv2-50M, and NTv2-250M (whose context windows are far shorter than these 84.6 to 409 kb genomes) were evaluated over overlapping fixed-length chunks. When one original sequence is represented by several chunks, per-chunk scoring is not directly comparable to per-sequence scoring. A single genome can contribute multiple predictions, changing the effective sample size and weighting classes by length. A cleaner protocol aggregates chunk-level outputs into one prediction per original sequence before computing metrics, for example by mean- or max-pooling the chunk embeddings, or by taking the majority vote or the mean of chunk-level NN scores (prior GFM detectors such as ViraLM likewise average per-fragment scores into a single sequence-level call [13]). Both the choice of aggregation rule and the number of retained PCA components are hyperparameters and must be selected on a dedicated validation split, kept strictly separate from the test set, to avoid optimistic leakage. In the present study, these choices were not tuned on an independent split, which is a limitation.

Second, the corruption experiments used random single-base substitutions and N-masking. These are convenient but only loosely mimic real sequencing errors and sample-preparation artefacts, which are non-uniform and technology-specific. Illumina data are dominated by substitutions, whereas nanopore reads are indel-rich, and library preparation introduces adapters, chimeras, and coverage bias. A more realistic robustness benchmark would therefore incorporate insertions and deletions and technology-specific error profiles (e.g., simulated with established read simulators). A recent multi-class oncoviral GFM classifier that tested both substitutions and indels found the models comparatively robust to substitutions but markedly more sensitive to indels than to substitutions, and the effect was reported for both tokenisation families [4]. For the NT, it is attributed to non-overlapping k-mer tokenisation, in which a single-base shift moves the reading frame, and comparable indel sensitivity is reported for the byte-pair tokenisation of DNABERT-S. Substitution-only corruption therefore understates real-world fragility, which applies to our own mutation and masking grid. Training-free perturbation probes, as introduced for recent generative DNA models [12], offer one template for such evaluation.

### Other limitations

Real metagenomic samples are imbalanced, fragmented, and may contain organisms absent from any reference. Sequences were sampled randomly rather than partitioned by taxonomy or collection date, and although C-RVDB is clustered at 98% identity, that step removes only near-identical records, so a query and its nearest indexed neighbour may still be almost identical. This matters more for retrieval than for supervised classification. Retrieval benchmarks do not normally draw queries from the indexed collection by random splitting, and the accepted practice for sequence data is to separate reference and query sets by identity or by taxonomy, for example through sequence clustering or genus-disjoint and temporal splits, under which foundation models are known to degrade substantially [16]. Our split therefore measures retrieval of near relatives, which is the regime in which alignment is inherently strongest, and the margin reported for BLAST should be read as a property of that regime as much as of the methods. The open-set setting in which GFMs are expected to excel, namely novel or highly divergent viruses with no close database homolog, was not directly tested and remains the key open question.

A related point concerns pre-training data. Corpora of this scale almost certainly contain viral sequences, and RVDB is itself assembled from the same public archives, so some overlap with our test data is to be expected. That overlap is not equivalent to an advantage. A pre-trained model has no mechanism for privileging those particular records. They are a vanishing fraction of a corpus spanning hundreds of species, the training objective never asked the model to tell them apart, and any individual organism is effectively diluted in the general representation that results. A custom BLAST database is the opposite case. It is restricted by construction to exactly the universe of organisms being classified, and every stored sequence is available for direct comparison. The reference collection therefore works for BLAST in a way that mere exposure during pre-training does not work for a foundation model, which is the same difference in mechanism discussed above.

### External evidence supports this expectation

When test contigs dissimilar to the training database are retained via a BLASTn similarity cut-off, FM-based identifiers lose only about 2% accuracy against about 10% for conventional convolutional and recurrent models [14].

### Future directions

Two NTv3 variants stand out. NTv3-8M, though weaker without adaptation, was the fastest model by a wide margin and its small parameter count suits task-specific fine-tuning under tight compute or hardware constraints, a promising basis for lightweight clinical tools. NTv3-650M achieved the best accuracy with competitive runtime and a highly compressible embedding space, making it the strongest candidate for large-scale deployment, with the main challenge being the compute and regularisation required to fine-tune 650M parameters on small viral datasets without overfitting.

Parameter-efficient fine-tuning offers a practical route around this. Low-rank adaptation has been shown to improve FM performance on downstream biological tasks while updating only a small fraction of parameters [32], and would allow the largest models to be specialised on viral data at modest cost. A complementary direction is multi-representation fusion, combining FM embeddings with explicit sequence descriptors such as k-mer spectra or physicochemical features, which has improved both viral identification [14] and protein-property prediction [32] over embedding-only baselines, and could also be extended to a hybrid retrieval pipeline that combines embedding similarity with alignment scores. Finally, evaluation on real metagenomic samples with orthogonally validated ground truth (e.g., PCR or culture confirmation) would substantially strengthen practical relevance, as would an explicit open-set benchmark probing generalisation to viruses held out of the reference under genus-disjoint or temporal splits [16].

## 5. Conclusion

We analyzed nine genomic foundation models for viral sequence classification from frozen embeddings, without any fine-tuning, on an RVDB-derived dataset, using FAISS nearest-neighbour search and BLAST as the alignment baseline. On sequences of standard length, BLAST (blastn, 0.97) outperformed every foundation model. On long sequences the two methods were closer, with NTv3-650M reaching 0.95 against 0.98 for blastn, a difference of about 0.02 that corresponds to six of the 256 test sequences and is not statistically established. PCA showed the NTv3-650M embedding space to be highly compressible, retaining more than 0.92 accuracy at 25% of its dimension. All foundation models were markedly more sensitive to base masking than BLAST. In summary, GFMs can already approach alignment-based methods on this task without any fine-tuning. NTv3-650M emerged as the most promising candidate for further optimisation and deployment, particularly for the open set and long-genome settings where alignment-based retrieval is expected to break down, while the much smaller NTv3-8M is the more attractive starting point where compute is constrained, being the fastest model by a wide margin and cheap enough to fine-tune.

## Acknowledgements

This work was supported by the project FloxgenAI: Platform for the Secure and Standardized Deployment of Foundation Models in Human Genome Analysis (CZ.01.01.01/01/24_062/0007508) and by the project Application of Genomic Transformer Models for Recognition of Patterns in RNA Fusion Sequences (SGS25/188/OHK4/3T/17).

## Competing interests

The authors declare no competing interests.

## Data and code availability

The analysis code, comprising Jupyter notebooks for dataset construction, embedding generation, nearest-neighbour classification, and BLAST evaluation, is available at https://gitlab.fel.cvut.cz/klempond/gfm-viral-identification. The Reference Viral Database (C-RVDB v31.0) is publicly available at https://rvdb.dbi.udel.edu/.

## Declaration of Generative AI and AI-assisted Technologies in Scientific Writing

During the preparation of this work, the authors used AI to improve the readability and language of the manuscript. After using this tool, the authors reviewed and edited the content as needed and take full responsibility for the content of the article. After refining the manuscript’s language with AI, it was further reviewed and enhanced to ensure clarity and accuracy.

## Notes

### Competing Interest Statement

The authors have declared no competing interest.

## References

1. Levine KS, Leonard HL, Blauwendraat C, et al. Virus exposure and neurodegenerative disease risk across national biobanks. Neuron. 2023;111(7):1086–1093.e2. doi:10.1016/j.neuron.2022.12.029.

2. Bedarf JR, Hildebrand F, Coelho LP, et al. Functional implications of microbial and viral gut metagenome changes in early stage L-DOPA-naïve Parkinson’s disease patients. Genome Med. 2017;9:39. doi:10.1186/s13073-017-0428-y.

3. Park SJ, Özdinç BE, Coker KG, Walsh DM, Fox DJ, Evans S, et al. Metagenomics indicates an interplay of the microbiome and functional pathways in Parkinson’s disease. npj Parkinsons Dis. 2026;12:60. doi:10.1038/s41531-026-01271-5.

4. Robertson J, Consul S, Vikalo H. NextVir: enabling classification of tumor-causing viruses with genomic foundation models. PLoS Comput Biol. 2025;21(8):e1013360. doi:10.1371/journal.pcbi.1013360.

5. Keeney JG, Gulzar N, Baker JB, Klempir O, Hannigan GD, Bitton DA, Maritz JM, King CHS IV, Patel JA, Duncan P, Mazumder R. Communicating computational workflows in a regulatory environment. Drug Discov Today. 2024;29(3):103884. doi:10.1016/j.drudis.2024.103884

6. Cassedy A, Parle-McDermott A, O’Kennedy R. Virus detection: a review of the current and emerging molecular and immunological methods. Front Mol Biosci. 2021;8:637559. doi:10.3389/fmolb.2021.637559.

7. Chin P-J, Bhavsar JD, Bosma TJ, MacDonald ML, Polson SW, Khan AS. Refinement of the Reference Viral Database (RVDB) for improving bioinformatics analysis of virus detection by high-throughput sequencing (HTS). mSphere. 2025;10(7):e00286–25. doi:10.1128/msphere.00286-25.

8. Altschul SF, Gish W, Miller W, Myers EW, Lipman DJ. Basic local alignment search tool. J Mol Biol. 1990;215(3):403–410.

9. Dalla-Torre H, Gonzalez L, Mendoza-Revilla J, Lopez Carranza N, Grzywaczewski AH, Oteri F, et al. Nucleotide Transformer: building and evaluating robust foundation models for human genomics. Nat Methods. 2024;22:287–297. doi:10.1038/s41592-024-02523-z.

10. Zhou Z, Ji Y, Li W, Dutta P, Davuluri R, Liu H. DNABERT-2: efficient foundation model and benchmark for multi-species genome. arXiv. 2023. arXiv:2306.15006.

11. Brixi G, Durrant MG, Ku J, Naghipourfar M, Poli M, Sun G, et al. Genome modelling and design across all domains of life with Evo 2. Nature. 2026;652(8112):1349–1361. doi:10.1038/s41586-026-10176-5.

12. Ben Allal L, Li Q, Fiusco M, Tunstall L, Rasul K, Beeching E, et al. Carbon: Decoding the Language of Life. bioRxiv. 2026. doi:10.64898/2026.05.22.727119.

13. Peng C, Shang J, Guan J, Wang D, Sun Y. ViraLM: empowering virus discovery through the genome foundation model. Bioinformatics. 2024;40(12):btae704. doi:10.1093/bioinformatics/btae704.

14. Wang Z, Yu Q, Li Y. VIRALpre: Genomic Foundation Model Embedding Fused with K-mer Feature for Virus Identification. bioRxiv. 2024. doi:10.1101/2024.11.12.623150.

15. Feng H, Wu L, Zhao B, Huff C, Zhang J, Wu J, et al. Benchmarking DNA foundation models for genomic sequence classification. bioRxiv. 2024. doi:10.1101/2024.08.16.608288.

16. Ye D, Hu F, Hu H, Hu S, Tan Y, Ouyang W, Li SZ, Cui J, Dong N. ViroBench: Benchmarking Nucleotide Foundation Models on Viral Genomics Tasks. Proc. 32nd ACM SIGKDD Conf. (KDD). 2026. arXiv:2605.25388. doi:10.1145/3770855.3819057.

17. Zhou Z, Wu W, Ho H, Wang J, Shi L, Davuluri RV, et al. DNABERT-S: learning species-aware DNA embedding with genome foundation models. arXiv. 2024. arXiv:2402.08777.

18. InstaDeep. Nucleotide Transformer v3 (NTv3): a foundational model for joint sequence and function. Technical report. December 2025. https://instadeep.com/wp-content/uploads/2025/12/NT_v3.pdf. Model cards: https://huggingface.co/InstaDeepAI

19. Johnson J, Douze M, Jégou H. Billion-scale similarity search with GPUs (FAISS). IEEE Trans Big Data. 2019.

20. Devlin J, Chang M-W, Lee K, Toutanova K. BERT: pre-training of deep bidirectional transformers for language understanding. NAACL. 2019.

21. Feng H, Wu L, Zhao B, Huff C, Zhang J, Wu J, et al. Benchmarking DNA foundation models for genomic and genetic tasks. Nat Commun. 2025. doi:10.1038/s41467-025-65823-8

22. Reimers N, Gurevych I. Sentence-BERT: sentence embeddings using Siamese BERT-networks. In: Proceedings of EMNLP-IJCNLP. 2019:3982–3992.

23. Li B, Zhou H, He J, Wang M, Yang Y, Li L. On the sentence embeddings from pre-trained language models. In: Proceedings of EMNLP. 2020. arXiv:2011.05864.

24. Walker PJ, Siddell SG, Lefkowitz EJ, et al. Changes to virus taxonomy and to the International Code of Virus Classification and Nomenclature ratified by the International Committee on Taxonomy of Viruses (2021). Arch Virol. 2021;166:2633–2648. doi:10.1007/s00705-021-05156-1

25. Simmonds P. A critique of the use of species and below-species taxonomic terms for viruses. Virus Evol. 2024;10:veae096. doi:10.1093/ve/veae096

26. International Committee on Taxonomy of Viruses. Master Species List MSL41 (2025-2026 release). 2026. doi:10.5281/zenodo.19154110

27. Smith DB, Bukh J, Kuiken C, Muerhoff AS, Rice CM, Stapleton JT, Simmonds P. Expanded classification of hepatitis C virus into 7 genotypes and 67 subtypes: updated criteria and genotype assignment web resource. Hepatology. 2014;59:318–327.

28. Matthijnssens J, Otto PH, Ciarlet M, Desselberger U, Van Ranst M, Johne R. VP6-sequence-based cutoff values as a criterion for rotavirus species demarcation. Arch Virol. 2012. doi:10.1007/s00705-012-1273-3

29. Fahmy AM, Ayad M, Ahmed HM. A unified benchmark of supervised and retrieval-based methods for viral genomic sequence classification. Sci Rep. 2026;16:27647. doi:10.1038/s41598-026-67272-9

30. Ghorbani A, Rostami M, Guzzi PH. AI-enabled pipeline for virus detection, validation, and SNP discovery from next-generation sequencing data. Front Genet. 2024;15:1492752. doi:10.3389/fgene.2024.1492752.

31. Wu L-Y, Wijesekara Y, Piedade GJ, Pappas N, Brussaard CPD, Dutilh BE. Benchmarking bioinformatic virus identification tools using real-world metagenomic data across biomes. Genome Biol. 2024;25:97. doi:10.1186/s13059-024-03236-4.

32. Infante S, Singh A, Kabir A. LoMuS: Low-Rank Adaptation with Sequence Multi-representation Improves Protein Stability Prediction. Bioinformatics. 2026;btag509. doi:10.1093/bioinformatics/btag509.

